# Numerical Investigation of Drug Transport from Blood Vessels to Tumor Tissue Using a Tumor-Vasculature-on-a-Chip

**DOI:** 10.1101/652099

**Authors:** Wei Li, Hao-Fei Wang, Zhi-Yong Li, Tong Wang, Chun-Xia Zhao

## Abstract

The delivery of adequate concentration of anticancer drugs to tumor site is critical to achieve effective therapeutic treatment, but it is challenging to experimentally observe drug transport and investigate the spatial distribution of the drug in tumor microenvironment. In this study, we investigated the drug transport from a blood vessel to tumor tissue, and explored the effect of tumor size, tumor numbers and positioning on drug concentration distribution using a numerical method in combination with a microfluidic Tumor-Vasculature-on-a-Chip (TVOC) model. The TVOC model is composed of a vessel channel, a tumor channel sandwiched with a porous membrane. A species transport model based on computational fluid dynamics was adapted to investigate drug transport. The numerical simulation was firstly validated using experimental data, and then used to analyse the spatial-temporal structure of the flow, and to investigate the effect of tumor size and positioning on drug transport and drug concentration heterogeneity. We found the drug concentration surrounding the tumor is highly heterogeneous, with the most downstream point the most difficult for drugs to transport and the nearest point to the blood vessel the easiest. Moreover, tumor size and positioning contribute significantly to this drug concentration heterogeneity on tumor surface, which is dramatically augmented in large and downstream-positioned tumors. These studies established the relationship between solid tumor size/positioning and drug concentration heterogeneity in the tumor microenvironment, which could help to understand heterogenous drug distribution in tumor microenvironment.

## 1. Introduction

Effective cancer treatment relies on the successful delivery of adequate drugs via the bloodstream to the tumor site, and the delivery of drugs to the tumor site correlates strongly with the structure of the vasculature and the properties of the tumor microenvironment. Tumor vasculature often has abnormal structures with leaky endothelial cell junctions thus hyperpermeability (*1, 2*). The hyperpeameability of tumor vasculature to nanoparticles or macromolecules is known as the enhanced permeability and retention (EPR) effect. Also, the tumor microenvironment is heterogenous with varied tissue components consisting of extracellular matrix, stromal cells, tumor cells, and immune cells. Consequently, these properties often lead to uneven drug distribution in tumor microenvironments (*3–5*) resulting in compromised treatment efficacy or even drug resistance which remains one of the major causes of treatment failure in clinics (*6–9*). However, drug transport from bloodstream to solid tumors has received very little attention.

Although it is known that drug extravasation is directly correlated with the fluid dynamics of blood flow (*10*), it remains largely unknown how drug transport in tumor microenvironment affects the spatial heterogeneity surrounding tumor tissue, which is composed of the aggressive and malignant layer of cancer cells (*11*). This is mainly due to the difficulty in determining drug distribution in tumor microenvironment *in vivo*. Microfluidics, as a powerful tool capable of precisely controlling and manipulating fluid flow, has been used to create *in vitro* systems that mimic *in vivo* drug extravasation, where drug distribution can be directly observed and analysed at a time scale of minute with multiple parameters independently controlled at a relatively reasonable cost (*12–18*). However, it remains challenging to obtain a complete dynamic map of drug concentration distribution due to the difficulties in high spatial- and temporal-resolution measurements. In contrast, computational fluid dynamics (CFD) enables researchers to conduct ‘numerical experiments’ to obtain detailed flow and drug distribution information.

In our previous study, we developed a drug extravasation model by implementing CFD with a two-phase model to analyse time-lapse drug accumulation, and demonstrated the effects of blood flow velocity, drug concentration and vascular permeability on drug concentration at the tumor channel (*10*), where the effect of tumors on drug transport were not taken into consideration. In this study, we constructed a Tumor-Vasculature-on-a-Chip model with the top layer mimicking the leaky blood vessel and the bottom channel containing different numbers of tumor spheroids with different sizes at different positions, and investigated the effect of tumors on drug transport with a focus on exploring drug concentration distribution in tumor microenvironment especially drug concentration heterogeneity around tumor periphery using a species transport model. This species transport model has been extensively employed to model convection and diffusion processes of multiphase mass transfer in environmental science (*19, 20*), energy industry (*21, 22*) and engineering (*23, 24*). It has also been adopted to study drug distribution in the eye (*25–27*), brain (*28*), lung (*29*), tumor nodule (*30*) and artery (*31*). We validated our numerical model with experimental data obtained using a TVOC device. Then we introduced a solid tumor with fixed size and position to the tumor channel, and studied the time-lapse drug distribution surrounding the tumor. We next varied either size or location of the tumor, and analysed the drug concentration heterogeneity on four different locations at the tumor surface. We also investigated the effect of multi tumors on drug concentration distribution. Lastly, we analysed the combined effect of tumor size and position on the drug concentration heterogeneity on tumor surface. To the best of our knowledge, this is the first study addressing drug transport and drug concentration heterogeneity using CFD.

## 2. Numerical Framework

### 2.1 Control equations for species transport model and fluid

Drug extravasation *in vivo* from tumor blood vessel surrounding tumor tissue is illustrated in Fig. 1(a). A microfluidic device was designed to mimic *in vivo* tumor microenvironment consisting of an upper channel mimicking the leaky tumor blood vessel and a bottom channel representing the tumor channel (Fig. 1(b)). A porous membrane sandwiched between the top and bottom channels has ten holes with the diameter of 8 µm to mimic the leaking vessel walls, and it has a membrane porosity of 0.02, which is defined as the ratio of the void hole areas to the total membrane areas. The blood vessel channel and tumor channel have a height of 270 µm, and length of 4000 µm and 7000 µm, respectively.

**Fig. 1.**
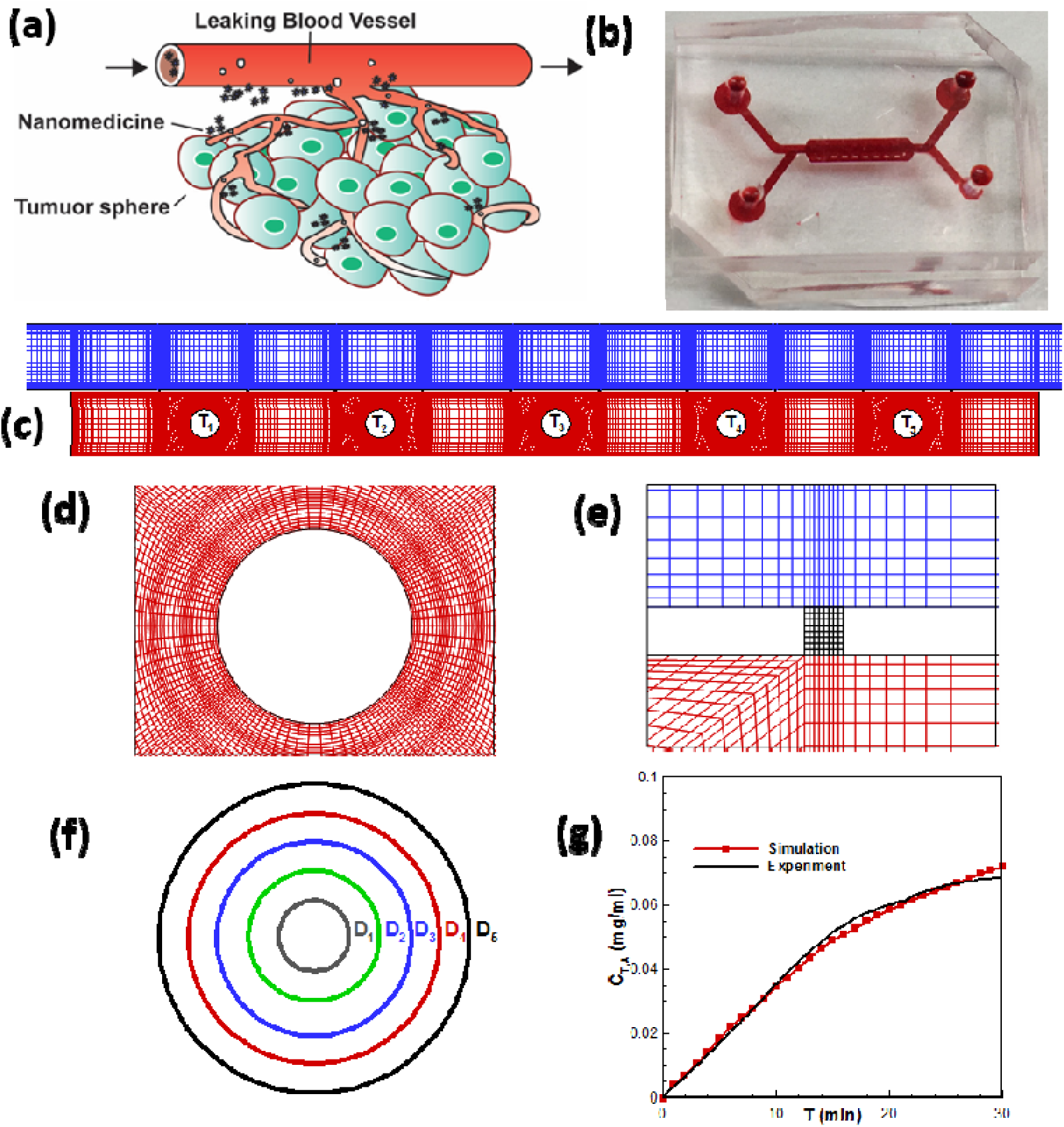
Microfluidic device setup and some pictures related to methodology. (a) Diagram of blood vessels near tumor; (b) Microfluidic device; (c) Computational mesh; (d) Mesh near tumor; (e) Mesh near the hole; (f) Five different tumor diameter sketch; (g) Comparison between experimental data and simulation results

The species transport model based on FLUENT 18.0 was adopted to simulate flow field in the top blood vessel channel and bottom tumor channel as well as drug transport from the top to the bottom channel. The microfluidic device was designed symmetrically. In addition, our preliminary steady-state 3D numerical study showed a typical 2D flow characteristic, a 2D model was therefore adopted. The unsteady two-dimensional Navier-Stokes governing equations were resolved using a finite volume method in order to reduce the computational resource.

The transport of drugs in the channels is governed by the species transport equation without considering the effect of temperature.

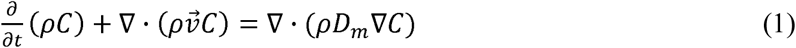

where C is the local concentration of the species, *D*_*m*_ is the mass diffusion coefficient for species in the mixture. In our case, there is no chemical reaction or any addition from the dispersed phase.

The solution of Equation (1) requires the velocity field and temperature field. The flow was modelled by pressure driven, single phase, laminar, incompressible flow and solutions were obtained by solving the Navier-Stokes Equations. The continuity equation, momentum equation with no body force terms, and energy equation with no heat sources are described as follows,

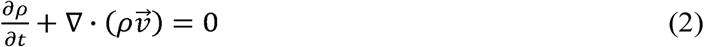

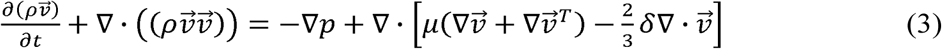

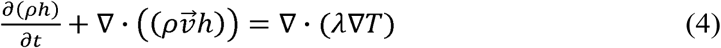

where 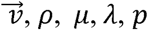, T and *h* are the velocity, density, molecule viscosity, effective conductivity, static pressure, static temperature, and internal energy.

### 2.2 Numerical method

The species transport model in FLUENT 18.0 was employed to conduct the numerical simulation of drug concentration distribution in the TVOC microfluidic device. We adopted the Finite-Volume Method (FVM) to solve the unsteady two-dimensional (2D) Navier–Stokes flow equations. The flow solution is water with a density of 998.2 kg/m^3^ and viscosity of 0.001003 kg/m·s. FITC-dextran was used as a model drug. The SIMPLE scheme was used to resolve the pressure-velocity coupling. The spatially discretization of the convection term was achieved by the second-order upwind scheme, so did the diffusion term with the second-order central difference scheme. The numerical investigation was implemented on high performance computer with ten nodes having double precision. For each case, firstly we obtained the steady converged result with the residual errors less than 10^−9^, then the unsteady calculation capable of investigating the drug extravasation process started with the steady converged result as its initial values. The time step was set to 0.1 second and it would undergo 20 iterations for each time step. To simulate drug concentration distribution at 30 min, it would work for 18000 time steps and take approximately 12 hours on high performance computer.

### 2.3 Computational mesh and boundary conditions

The computational mesh of the numerical simulation has a geometry comprising three distinct and connected parts (Fig. 1(c)). Blue meshes refer to the main channel, which represents the blood flow in normal blood vessel. Red meshes display the tumor channel. The mesh is generated in ANSYS ICEM. Five different tumor positions are marked as T_1_, T_2_, T_3_, T_4_, and T_5_ (Fig. 1(c)), each represents the positioning of a tumor with the diameter of 150 µm that distributed equally from the left side to the right side of the tumor channel, with T_3_ in the middle of tumor channel.

Figs. 1(d)-(e) display the mesh near the hole and the tumor spheriod. Fig. 1(f) shows the diameter sketches of five different diameters: D_1_ (50 µm), D_2_ (100 µm), D_3_ (150 µm), D_4_ (200 µm), and D_5_ (250 µm). All meshes are structural, and mesh independence test was performed to ensure the convergence of the numerical simulation. Two mesh sizes with their numbers of approximate 60,000 and 120,000 were tested, respectively. As they showed no difference in the area-weighted average spatial-temporal drug concentration, we therefore selected the 60,000-noded mesh for this investigation. In addition, grid independence was maintained to specifically compare the changes induced by the variation in tumor size and positioning.

The inlet velocity and outlet static pressure boundary conditions are adopted and the non-slip and adiabatic wall boundary are employed at the walls. For the inlet flow, the uniform inlet velocity is set to 0.3 mm/s to mimic the averaged blood flow velocity in capillary (*32*), with inlet total temperature of 300 K, which represents the actual velocity of the human blood flow. The initial pressure is set to 101,330 Pa with uniform inlet drug concentration specified to 0.10 mg/ml.

### 2.4 Validation of the numerical method

Experimental methods were described in our previous study (*10*). Based on the experimental facility, we conducted the drug transport experiment. The comparison between the time-lapse curve of the simulated and experimental data of area-weighted average drug concentration (C_T,A_) over the tumor channel are shown in Fig. 1(g). Area-weighted average drug concentration (red line) firstly shows a linear growth, then slows down and eventually reaches the near-plateau equilibrium state, due to the limit of the channel volume. The simulation data (black line) fits well with the experimental data, thus validating our numerical methodology (Fig. 1g).

## 3. Results

### 3.1 Effect of tumor size on drug concentration distribution in the tumor channel

To investigate the effect of tumor size on drug transport, we compared the average drug concentration in the tumor channel with tumor embedded. Five different tumor diameters (D_1_ to D_5_) including 50, 100, 150, 200, and 250 μm, were studied. The tumor spheroid is located in the middle of the tumor channel (T_3_). Two different types of drug concentrations were defined here to differentiate average drug concentrations of the whole tumor channel (C_T,A_) versus local drug concentrations (C_T_) of a specific point on the tumor spheroid surface in the tumor channel. Fig. 2(a) shows the time-lapse average drug concentration in the tumor channel with tumors having five different tumor sizes (D_1_ to D_5_). We can see that the tumor size does not have significant effect until it is bigger than 200 µm which is close to the depth of the tumor channel (270 µm). The average drug concentration in the tumor channel having a tumor of 250 µm in diameter is much smaller than those with smaller tumor sizes over the 30 min time course, demonstrating the blocking effect of bigger tumors. Although the average drug concentration increases with time for all different tumor sizes, its growth rate decreases as the tumor diameter grows. Especially, with the tumor diameter exceeding 200 µm, the average drug concentration drops dramatically. The comparison between average drug concentration values at 30 minute (C_T,A30_) shows little change (around 0.073 mg/ml) when the diameter varies from 50 to 200 µm (D_1_ to D_4_). However, C_T,A30_ drops to 0.048 mg/ml when the dimeter reaches 250 µm (D_5_), which is 34% of the average drug concentration for D_1_ to D_4_. These results demonstrates the effect of tumor size on drug transport from the vessel channel to the tumor channel, the bigger the tumor the lower drug concentration in the tumor channel.

**Fig. 2.**
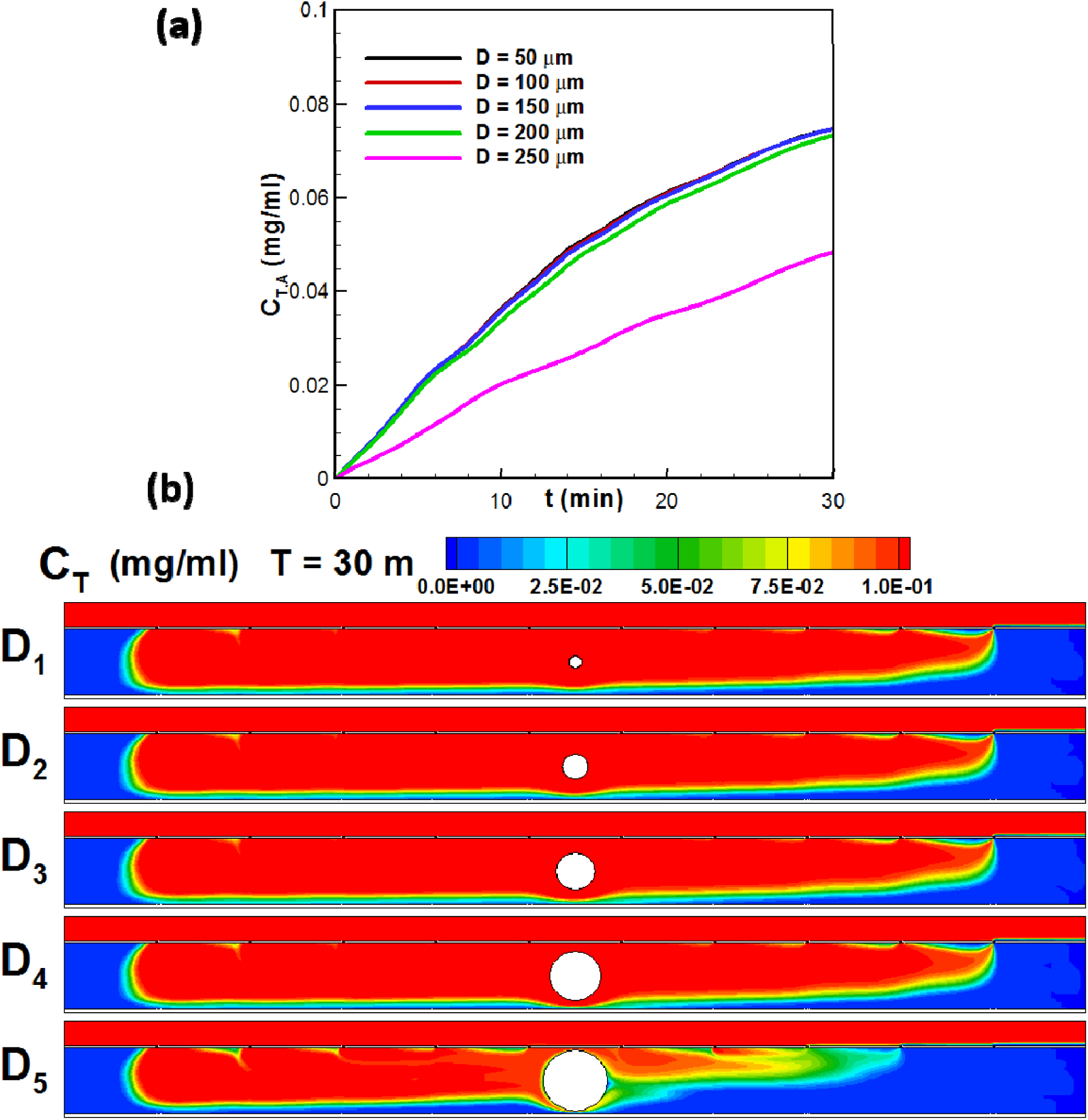
Drug concentration distribution in the tumor channel with five different tumor diameters. (a) Average drug concentration in the tumor channel vs time; (b) Local drug concentration contours at 30 min.

As local drug concentration adjacent to the tumor determines the uptake of the drug by tumor tissue (*33*), we further investigated the impact of tumor size on the local drug concentration at the tumor surface. Fig. 2(b) shows the spatial local drug concentration distribution at 30 min of tumors having diameters of D_1_ to D_5_ at T_3_ position of the tumor channel. We found that the distribution of local drug concentration in the tumor chamber is uneven. This non-uniform distribution is further aggravated by the increase of tumor diameter, and leads to the heterogeneity of local drug concentration encircling the tumor. To analyse the temporal changes of this heterogeneity, we analyzed the local drug concentrations of four points on the tumor surface, the leftmost (P_1_), rightmost (P_2_), bottom (P_3_), and top (P_4_) point.

For tumors with diameters of 50 µm (D_1_), 100 µm (D_2_), 150 µm (D_3_), 200 µm (D_4_) and 250 µm (D_5_), Figs. 3(a)-(e) show the temporal local drug concentrations at the four points (P_1_ - P_4_), and Fig. 3(f) illustrates the local drug concentrations at 30 min. As the tumor spheroid is located at the middle of the tumor channel (T_3_), it takes some time for the drug travelling from the inlet to the tumor, then the drug arrives at the tumor (Fig. s2), it quickly reaches the equilibrium for tumors of smaller sizes (50 – 200 µm). Therefore, the local drug concentration distribution demonstrates a slow growing stage followed by a rapid growing stage and finally a plateau stage. For the biggest tumor (250 µm), drug transport was much slower because of the blocking effect, therefore, equilibrium concentrations were not observed within the 30 min time scale.

**Fig. 3.**
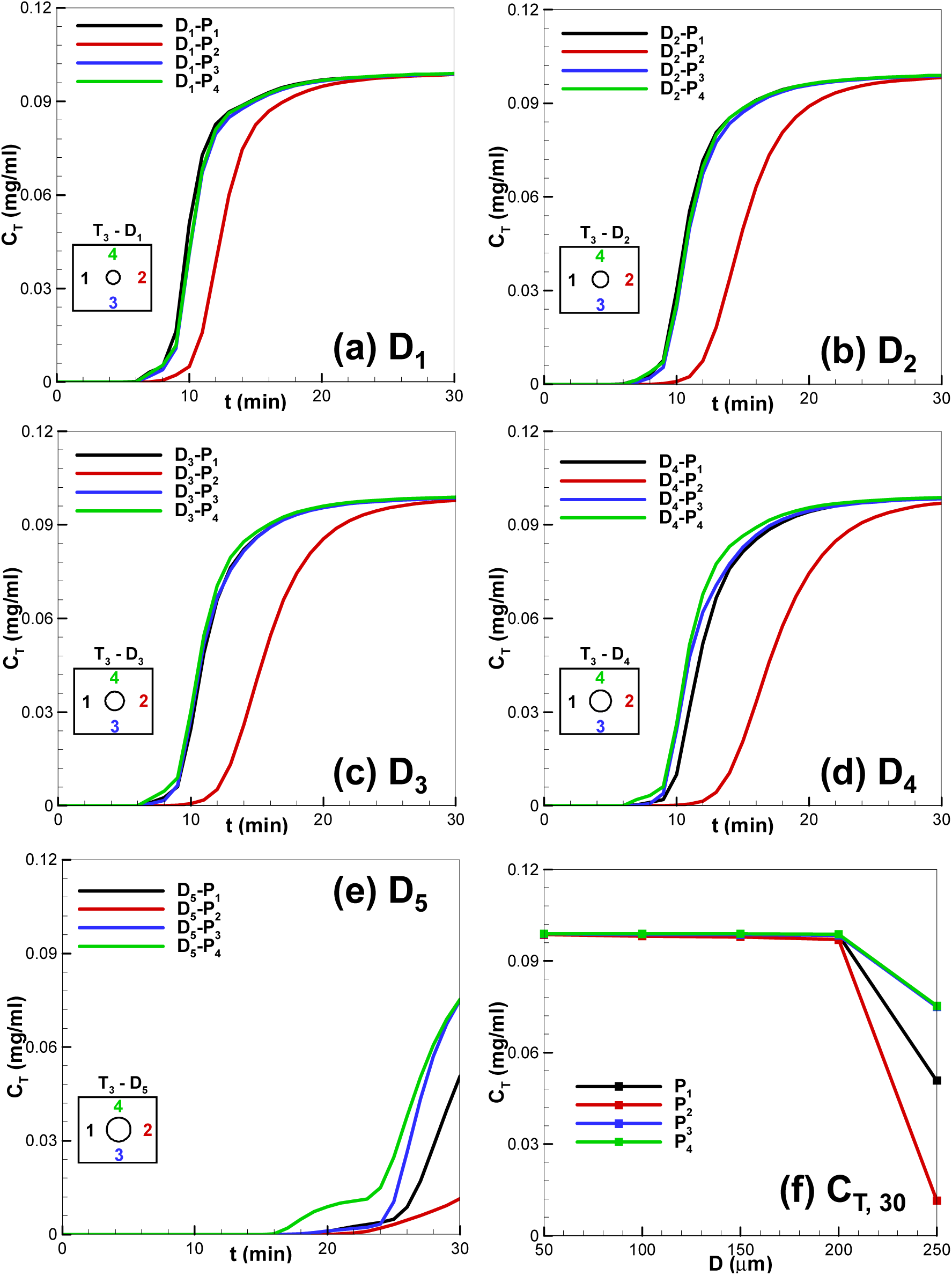
Local drug concentrations of four positions 1, 2, 3 and 4 at the tumor surface with five different tumor diameters. (a) D_1_; (b) D_2_; (c) D_3_; (d) D_4_; (e) D_5_; (f) Local drug concentrations of tumor spheroids with different sizes at the four points P_1_, P_2_, P_3_, and P_4_ at 30 min.

Interestingly, for tumors with all the five diameters, the local drug concentration at P_2_ is discrete from the rest points and dramatically delayed in reaching the equilibrium (Fig. 3(a)-(d)). Furthermore, a significantly lower local drug concentration (C_T30_) is observed when the diameter is larger than 200 µm (Fig. 3(f)). These kinetic results suggest the heterogeneous local drug concentration surrounding the tumor. The far most P_2_ position is the most difficult location and the top point P_4_ is the easiest location for drug to reach equilibrium. The much lower concentration at P_2_ is mainly caused by the adjacent recirculation around the tumor, preventing the flow of drug from arriving at the far-most tumor surface (Fig. s1(b)).

### 3.2 Impact of tumor positioning on drug concentration distribution

In addition to the size of the tumors, another critical factor, the relative tumor position in tumor channel, affects drug transport. Fig. 4 shows the drug concentration distribution at five different X-direction positions (T_1_ - T_5_), which are distributed equally from the upstream (left side) to downstream (right side) of the tumor channel, with T_3_ at the middle of the tumor channel. The results of two tumor diameters are presented, namely 150 µm (D_3_) and 250 µm (D_5_). Similar to Fig. 2(a), the average drug concentration increases as time lapses for all five different tumor positions (Fig. 4(a)). For the tumor with 150 µm diameter (D_3_), the average drug concentration distribution is not affected by the position of the tumor, which is reflected by the overlapping average drug concentration curves from T_1_ to T_5_. However, for the tumor size of 250 µm, the average drug concentration decreases significantly. The closer the tumor to the middle of channel the lower the average drug concentration is. It decreases gradually from T_1_ to T_3_, and it reaches the minimal value at T3; and then it gradually increased from T_3_ to T_5_. The average drug concentration at T_1_ is nearly the same as that at T_5_. Similarly, the concentration at T_2_ is almost the same as that at T_4_, mainly because the flow field is almost symmetrical with respect to the middle vertical line of the tumor channel (Fig. s1), so do the velocity and drag force on tumors (Fig. s3). The drag force on D_3_ tumor is smaller than that on D_5_ tumor, so the average drug concentration with D_3_ tumor is larger than that with D_5_ tumor. The drag force on D_5_ tumor in T_3_ location is the largest, so its average drug concentration is the minimum. In summary, the positioning of D_5_ tumor plays an important role in regulating drug concentration distribution.

**Fig. 4.**
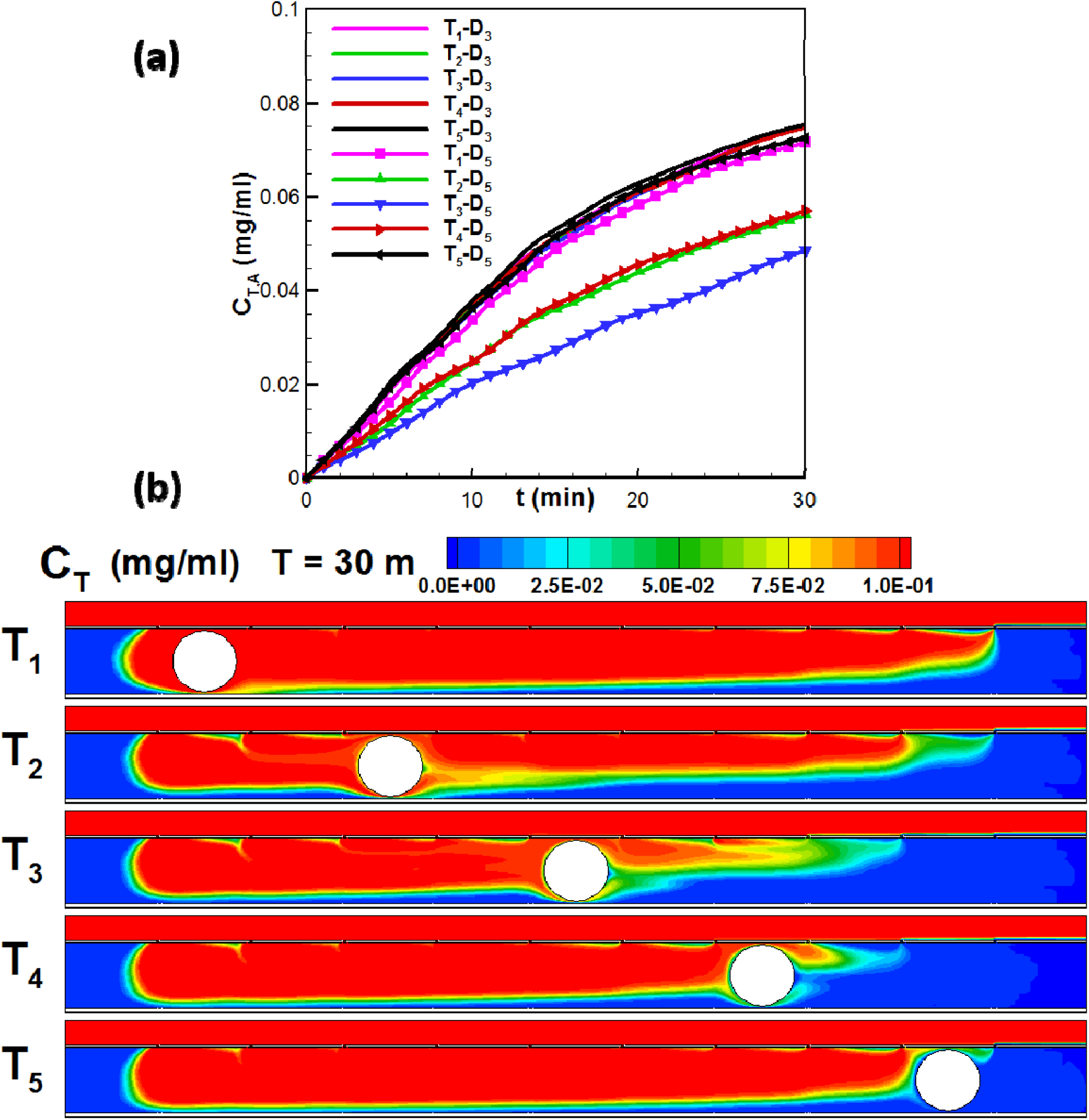
Drug concentration distribution with five different tumor locations. (a) Average drug concentration vs time; (b) Drug concentration contours at 30 min.

To better describe the impact of tumor positioning on the local drug concentration on the tumor surface, we investigated the spatial local drug concentration distribution for five different tumor locations (T_1_-T_5_) at the time of 30 min (Fig. 4(b)), with the tumor diameter fixed at 250 µm (D_5_). The closer the tumor positioning to the middle of channel the less the average drug concentration is. In addition, the more downstream the tumor is, the more difficult for the drug to transport to the tumor surface.

For tumors at positions T_1_ to T_5_, the temporal local drug concentration on the four different surface locations (P_1_ to P_4_) are shown in Fig. 5(a)-(e), with its distribution on 30 min are shown in Fig. 5(f). We fixed the tumor diameter at 150 µm (D_3_). Similar to Fig. 3, for the tumor with D_3_ diameter at T_1_ to T_4_ position, the local drug concentration change also consists of three stages: a slow growing stage, then a rapid growing stage and finally a plateau stage. However, for the T_5_ position, only the first two stages appear, indicating the local drug concentration fails to achieve the equilibrium at T_5_ position. For all locations, P_2_ is the most difficult point (short-board) and P_4_ is the easiest point (long-board) for drugs to reach. This further demonstrates that the tumor positioning has an important effect on the local drug concentration heterogeneity at tumor surface. Fig. 5(f) shows the local drug concentration on tumor surface at 30 min. The local drug concentration is significantly reduced for the T_5_ position, suggesting the downstream T_5_ is the most difficult position for drugs to reach.

**Fig. 5.**
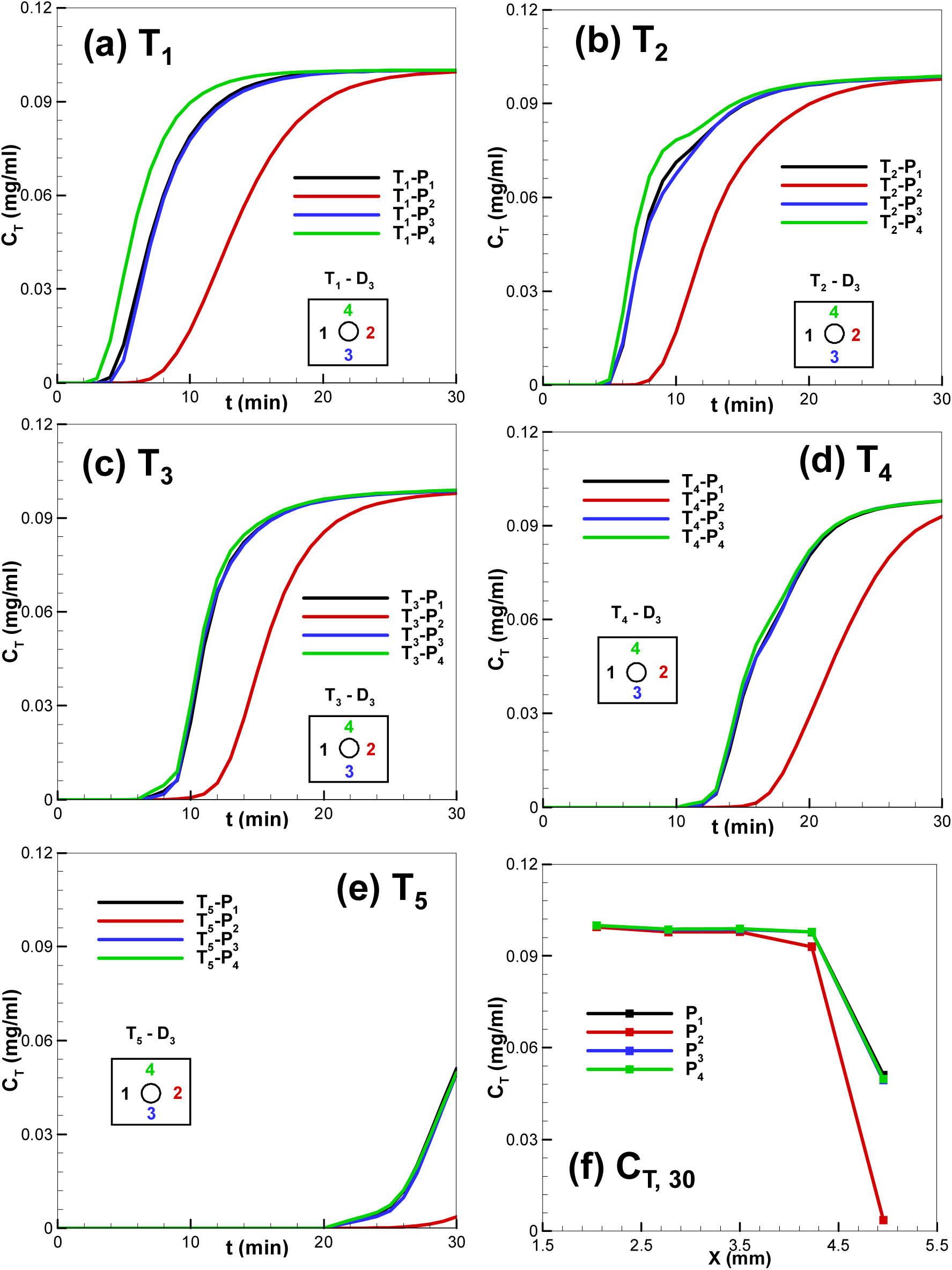
Drug concentrations of four positions 1, 2, 3 and 4 at the tumor surface with five different tumor locations. (a) T_1_; (b) T_2_; (c) T_3_; (d) T_4_; (e) T_5_; (f) Local drug concentrations at the four points P_1_, P_2_, P_3_, and P_4_ at different positions at 30 min.

### 3.3 Impact of multi-tumor on drug concentration distribution

To investigate the effect of multiple tumors on drug extravasation, we compared the drug concentration distribution with three or five tumors embedded. Fig. 6(a) shows the average drug concentration in the tumor channel with time. The lines with and without symbols refer to multiple tumors with 250 µm (D_5_) and 150 µm (D_3_), respectively. We can clearly see that the average drug concentration remains the same for multi tumors with diameters of 150 µm (D_3_). However, for tumors with diameters of 250 µm (D_5_), the increase of tumor numbers result in the decrease of average drug concentration. C_T,A_ with five tumors is smaller than that with three tumors. For three tumors in the channel, its C_T,A_ with tumors at T_2_ - T_4_ location is smaller than that at T_1_, T_3_, and T_5_, demonstrating that the tumor positioning has an important effect on the average drug concentration for tumors with bigger size but not small ones. Fig. 6(b) shows the spatial local drug concentration distribution contours with three or five D_5_ tumors at 30 min. Three tumors at T_2_ - T_4_ locations have more blockage effect on drugs than that in T_1_, T_3_, and T_5_ location. C_T,A_ with five tumors located at positions T_1_ – T_5_ just reduces a little relative to three tumors, because the tumor located at T_1_ or T_5_ just decrease the drug concentration slightly.

**Fig. 6.**
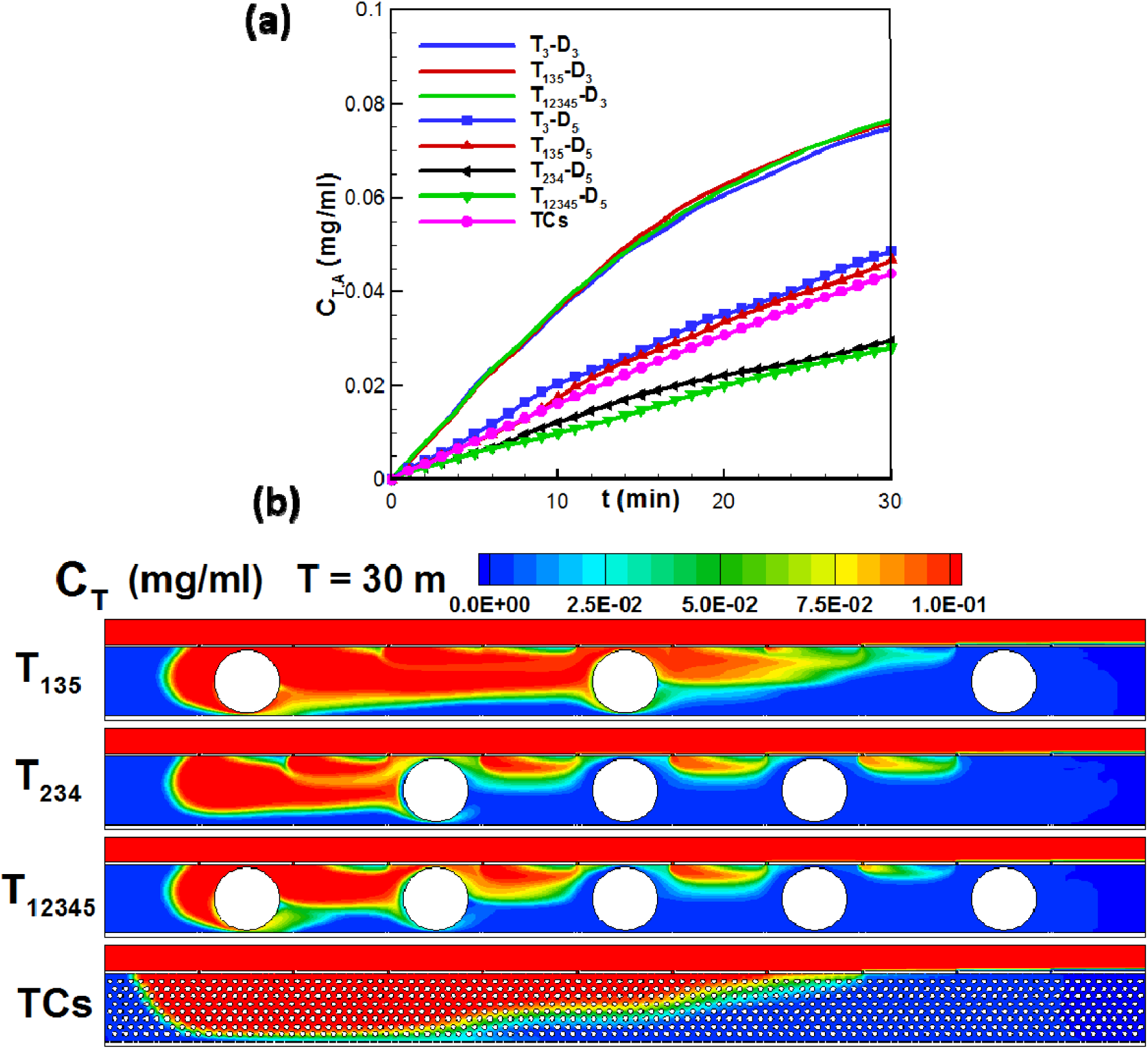
Drug concentration distribution in the tumor channel with multiple tumors.(a) Average drug concentration vs time (D_5_: lines with symbols; D_3_: lines; Pink line: multiple tumor cells); (b) Drug concentration contours at 30 min with D_5_ and multiple tumor cell aggregates.

In addition, we also investigate the drug transport in the tumor channel consisting of multiple small tumor cell aggregates. 796 tumor cell aggregates with the diameter of 20 µm are evenly distributed in the tumor channel. The pink line with circle symbols in Fig. 6(a) indicates its averaged drug concentration. Although the total area of tumor cell aggregates is more than five times of the tumor surface for the diameter of 250 µm (D_5_), its average drug concentration is higher than that with five tumors at positions T_2_ - T_4_, indicating it is easier for drugs to transport to most tumor cells relative to big five tumor spheroids with diameter of 250 µm (D_5_). The lowest contour in Fig. 6(b) displays the drug concentration at 30 min, where it is bigger than that with five tumors in T_1_ - T_5_ and three tumors in T_2_ - T_4_. This result indicates that the drug is still accessible for smaller tumors even so many tumor aggregates but less dense extracellular matrix. In contrast, much less drug can be delivered to tumor tissue of big sizes and dense structure.

### 3.4 Analysis of local drug concentration heterogeneity on tumor surface

Based on the above analysis, we found that among all the surface locations, the P_2_ and P_4_ are the most difficult and easiest location for drugs to diffuse to, respectively. Accordingly, the C_T_ value on P_2_ is the maximum and on P_4_ is the minimum. We next investigated the drug concentration heterogeneity on the tumor surface based on the local C_T_ value of P_2_ (short-board, Fig. 7(a)) and P_4_ (long-board, Fig. 7(b)) at 30 min for five different tumor locations (T_1_-T_5_) and five different tumor diameters (D_1_-D_5_). The x-axis and y-axis are the tumor locations (T_1_-T_5_) and tumor diameters (D_1_-D_5_), respectively. We found that the plateau C_T_ (red-domain) is achieved for tumors with diameters ranging from D_1_ to D_4_ and positioned from T_1_ to T_4_. However, when the tumor diameter exceeding 200 µm (D_4_) or positioned further than T_4_, the local drug concentration reduced sharply on the short-board P_2_ and eventually reached the lowest point (D_5_, T_5_). Similar trend was also observed for the long-board P_4_ (Fig. 7(b)), with a much slower decreasing rate and higher final drug concentration at the lowest point (D_5_, T_5_). We further analysed drug concentration heterogeneity using the local drug concentration difference ΔC_T_, which is defined as ΔC_T_ = C_T,P2_ C_T,P4._ As shown in Fig. 7(c), we found within the zone of (T4, D4), ΔC_T_ had very low values, whereas outside this zone, ΔC_T_ gradually increased. These results indicated that both the tumor size and its relative location greatly affect the drug heterogeneity on the tumor surface.

**Fig. 7.**
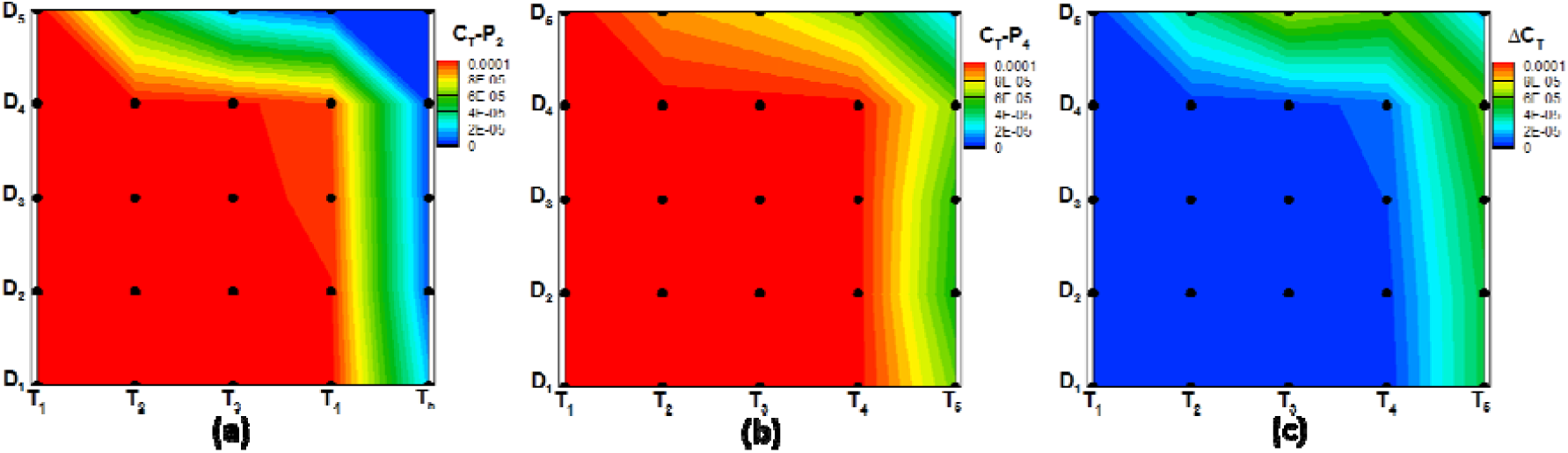
Effect of tumor positioning and locations on drug concentration heterogeneity on P_2_ and P_4_. (a) Local drug concentration on point P_2_; (b) Local drug concentration on point P_4_; (c) Local drug concentration difference between points P_2_ and P_4_.

## 4. Conclusion

In this study, we investigated numerically drug transport from blood vessel to tumor tissue using a Tumor-Vasculature-on-a-Chip model, and demonstrated that the drug concentration surrounding the tumor was highly heterogeneous, with the most downstream point the most difficult for drugs to arrive at. Then we further explored the effect of tumor size, positioning as well as the density of tumor tissues on drug transport, and found that solid big tumor tissues had significant blocking effect, the transport of drugs from the vessel channel to the tumor channel was significantly hindered. In contrast, the blocking effect of many small tumor cell aggregates was not as significant as the big tumor spheroids. Although our numerical simulation is based on a simplified model, these results provide useful information for understanding the transport barrier of drugs from tumor vasculature to tumor tissue. Future research is warranted to take into account of more *in vivo* factors such as the porosity of tumor extracellular matrix and stroma cells, which varies greatly in different types of cancers. Meanwhile, we will also study the properties of various drug carriers, such as their size, shape, hardness, affinity with tumor extracellular matrix to provide deeper insights into drug transport in tumor microenvironment.

## Acknowledgement

We acknowledge the supports from High Performance Computing of Queensland University of Technology. C.-X Zhao acknowledges financial support from the award of the Australian Research Council (ARC) Future Fellowship (FT140100726). TW acknowledges financial support from the award of the Australian Research Council DECRA Fellow (DE170100546). This work was performed in part at the Queensland node of the Australian National Fabrication Facility (ANFF-Q), a company established under the National Collaborative Research Infrastructure Strategy to provide nano and micro-fabrication facilities for Australia’s researchers.

