## Supplemental materials for "Numerical Investigation of Drug Transport from Blood Vessels to Tumor Tissue Using a Tumor-Vasculature-on-a-Chip"

**Supplementary Information**

**Flow structure analysis**

The flow field distribution was simulated using the above described numerical methods. The velocity of inlet flow is 0.3 mm/s. The membrane porosity of blood wall is set at 0.02. The tumor position is the third one in the middle of channel and its diameter is 150 µm. Only the flow field at the time of 30 min was analysed, as the blood flow was nearly steady even in the unsteady simulation. Fig. s1(a) shows the velocity field in both channels, with the laminar boundary layer boundary fully established just over the tumor channel. Fig. s1(b) displays the velocity distribution of the tumor channel, which is much smaller than that of vessel channel. The flow velocity below holes is big due to the formation of jets there. Overall, the flow velocity progressively increases from the inlet to the middle of the tumor channel. This is due to the constant entry of the main flow to the tumor channel from H_1_ to H_5_, and then the flow velocity reduces gradually as it passes through H_6_ to H_10_. Similar to the flow surrounding a cylinder, the velocity upstream of the tumor is smaller due to blockage by the tumor. Velocity downstream of the tumor is also reduced because the low velocity zone appears. Fig. s1(c) shows in both channels, the distribution of Y-velocity with formation of jets at holes. It can be observed that the drug flow enters the tumor channel through H_1_ to H_5_. The Y-velocity peaks at H_1_ then decreases from H_1_ to H_5_. The flow exit through H_6_ to H_10_, where the Y-velocity is the largest for the last hole (H_10_) and then gradually reduces from the last hole (H_10_) to the last four hole (H_9_-H_6_). In addition, the direction of Y-velocity along tumor changes when the flow moves along the tumor. Fig. s1(d) shows the streamline distribution in tumor channel, where 2 vortexes are formed near both ends, which explains the low velocities near both ends in Fig. s1(b). The streamline from H_1_ stems from the H_10_. Similarly, the streamline from H_2_ stems from H_9_, and so on so forth, until H_5_ to H_6_.

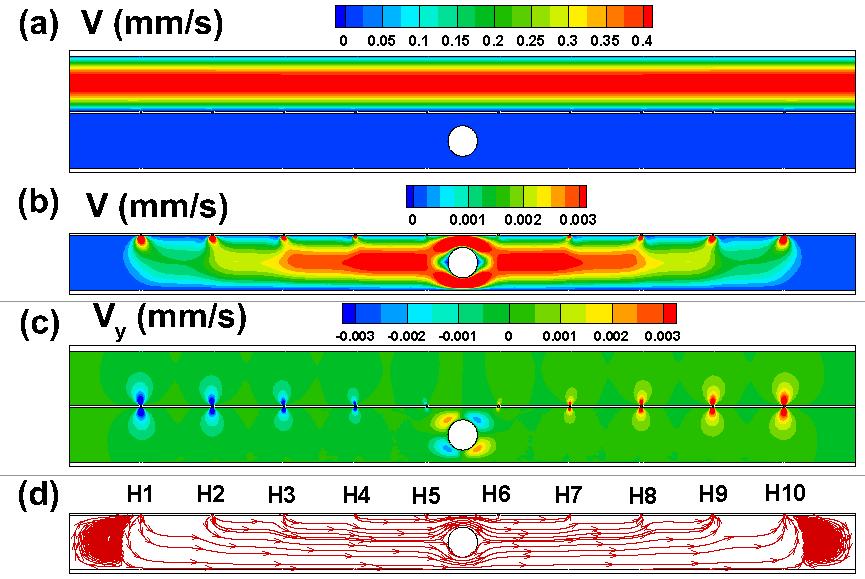

Fig. s1 Flow field analysis at 30 min. (a) Velocity distribution in both channels; (b) Velocity distribution in tumor channel; (c) Y-velocity distribution near the membrane; (d) Streamline distribution in tumor channel.

**Drug transport process**

Fig.s2 shows the drug concentration distribution in tumor channel at the time of 1, 10, 20, and 30 min to describe the drug transport process from the upper channel to the tumor channel. At the time of 1 min, the drug enters into tumor channel from the first five holes. In addition, drugs convected with blood flow do not arrive at the tumor surface, so the drug concentration on tumor surface increases slowly. With drugs start to convect to tumor surface at the time of about 10 min, the drug concentation on tumor surface increase rapidly till to the time of 20 min, in which drugs transported to tumor surface, indicating the drug concentration almost reach the equilibrium. During the period between 20 and 30 min, the drug concentration on tumor surface increase slowly to reach the plateau. In total, there are three stages of drug concentration increasing on tumor surface. The first is very slowly increasing stage followed by a rapid increasing stage, and then the plateau is gradually achieved at the final stage.

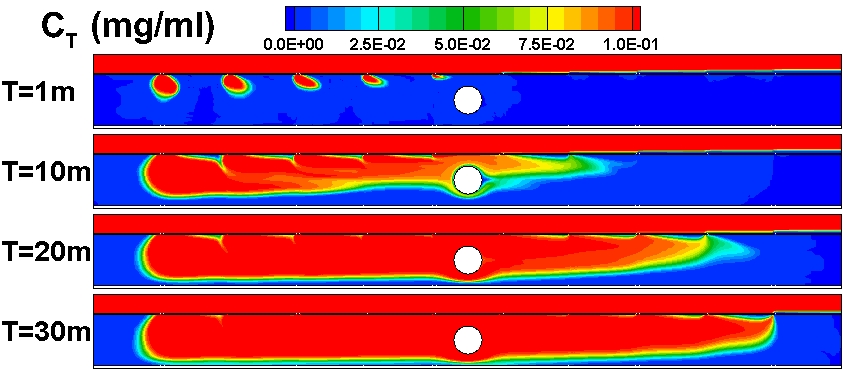

Fig.s2 Drug concentration distribution in tumor channel at 1, 10, 20, and 30 min

**Drag force on tumors and velocity distribution**

Fig. s3 displays the average velocity distribution and drag force on tumors. Fig. s3(a-b) shows the average velocity distribution along the X-direction in the tumor channel in 5 different tumor positions with the tumor diameter of 150 and 250 µm, respectively. The velocity gradually increased from the left side to the middle part of tumor channel, and then it started to decrease. The tumor blocked the flow and reduced the area the flow can pass through, so the velocity near the tumor increased. For the diameter of 250 µm, its fluctuation aptitude increased more than that of the tumor diameter of 150 µm. In addition, the velocity distributions are nearly symmetrical based on the middle vertical line. Fig. s3(c) is the drag force distribution on tumors in five different tumor positions. In total, the drag force with the diameter of 250 µm was larger than that with the diameter of 150 µm. It nearly remained the same with the diameter of 150 µm. For the diameter of 250 µm, it increased gradually from T_1_ to T_3_ and reached the maximum at T_3_, then it began to reduce. The increase of the drag force blocked the drug to enter into the tumor channel, so the drug concentration reduced.

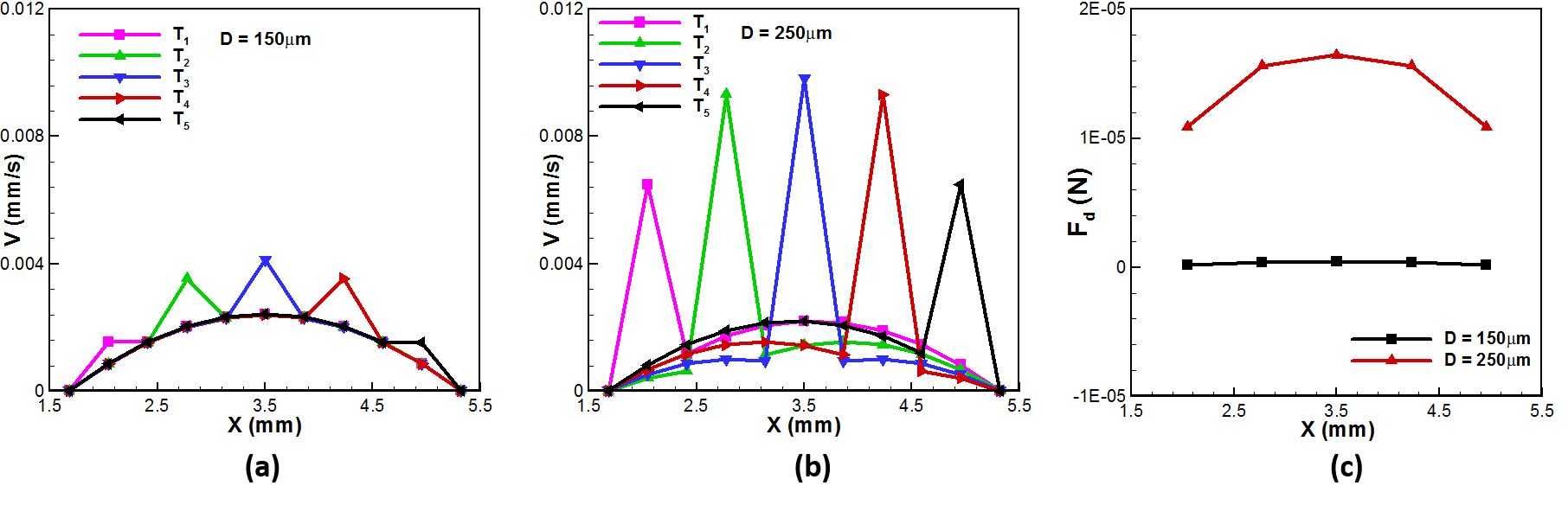

Fig. s3 Average velocity distribution and drag force on tumors. (a) Average velocity distribution along X-direction in tumor channel in 5 different tumor location with the tumor size of D_3_; (b) Average velocity distribution along X-direction in tumor channel in 5 different tumor location with the tumor size of D_5_; (c) Drag force on tumors in 5 different tumor locations with the tumor size of D_3_ and D_3_ , respectively.
